# Clinical and Functional Characterization of Melanocortin 4 Receptor genetic variants in African American and/or Hispanic children with severe early onset obesity

**DOI:** 10.1101/491969

**Authors:** Maria Caterina De Rosa, Alessandra Chesi, Shana McCormack, Justin Zhou, Benjamin Weaver, Molly McDonald, Sinead Christensen, Kalle Liimatta, Michael Rosenbaum, Hakon Hakonarson, Claudia A. Doege, Joel N. Hirschhorn, Struan F.A. Grant, Vidhu V. Thaker

**Affiliations:** Columbia University Medical Center, New York, NY 10032; Children’s Hospital of Philadelphia, Philadelphia, PA 19104; Boston University School of Medicine, Boston, MA 02118; Boston Children’s Hospital, Boston, MA 02115

## Abstract

**Context:** Mutations in melanocortin receptor (*MC4R*) are the most frequent cause of monogenic obesity in children of European ancestry, but little is known about their prevalence in children from minority populations in the United States.

**Objective:** This study aims to identify the prevalence of *MC4R* mutations in children with severe early onset obesity of African-American and/or Latina ancestry.

**Design and Setting:** Individuals were recruited from the weight management clinics at two hospitals and from the institutional biobank at a third hospital. Sequencing of the *MC4R* gene was performed by whole exome and/or Sanger sequencing. Functional testing was performed to establish the surface expression of the receptor and cAMP response to its cognate ligand α-melanocyte stimulating hormone.

**Participants:** Three hundred and twelve children (1-18 years, 50% girls) with body mass index (BMI) > 120% of 95^th^ percentile of CDC 2000 growth charts at an age < 6 years, with no known pathological cause of obesity were enrolled.

**Results:** Eight rare *MC4R* mutations (2.6%) were identified in this study (R7S, F202L (n=2), M215I, G252D, V253I, I269N, F284I), three of which have not been previously reported (M215I, G252D, F284I). The pathogenicity of the variants was confirmed either by prior literature reports, or by functional testing. There was no significant difference in the BMI or height trajectories of children with or without *MC4R* mutations in this cohort.

**Conclusions:** While the prevalence of *MC4R* mutations in this cohort was similar to that reported in obese children of European ancestry, some of the variants were novel.

## Introduction

The leptin melanocortin axis plays a crucial role in weight regulation, with mutations in gene members being identified as causal for human monogenic obesity(1). The melanocortin 4 receptor (MC4R), which is most abundantly expressed in the paraventricular nucleus of the hypothalamus, is the sensor for orexogenic, as well as anorexogenic, signals in this pathway(2). Binding of alpha-melanocyte stimulating hormone (α-MSH) to the G-protein coupled receptor MC4R induces satiety. Mutations in the *MC4R* gene and genes encoding its trafficking pathway have been identified as the most common cause of monogenic obesity in children of European ancestry, with prevalence estimates ranging from 1.5 to 5.8%, mostly comprised of mutations in *MC4R* (3–5). Individuals with *MC4R* mutations have been reported to have significant hyperphagia and early onset of obesity, even in the heterozygous state(3–5).

While the obesity epidemic has disproportionately affected non-Hispanic African-American and Hispanic youth in the United States (6), little is known about the prevalence of *MC4R* mutations in children of these ethnicities. In a recently published study of 25 unrelated Afro-Caribbean children with obesity from the Guadeloupe Island, one 11.8 year-old boy had a p. Ile301Thr mutation in *MC4R* (7), while in a study of South African adults of mixed ancestry that included 63% Black Africans (n = 187), 2.6% harbored a rare *MC4R* variant (8). These studies provide early clues on the expected prevalence of *MC4R* variants in children of non-European ancestry. In this study, we catalog the frequency of *MC4R* mutations in children with severe early onset obesity of African-American and/or Latina ancestry in the United States.

## Materials and Methods

### Clinical study recruitment

The study cohort was recruited from Boston Children’s Hospital (BCH), Children’s Hospital of Philadelphia (CHOP) and Columbia University Medical Center (CUMC), under each institution’s IRB approval. Informed consent, and where appropriate, assent was obtained prior to enrollment. The inclusion criteria were presence of severe early onset obesity, defined as body mass index (BMI) > 120% of the 95^th^ percentile on the CDC 2000 BMI charts for age and sex (BMI_95pct_) (9) documented under 6 years of age with no previously known underlying genetic or other pathogenic cause; for the sequencing study reported herein, we focused on children with selfreported African American and/or Latina ancestry with age at recruitment between 2-18 years. The subjects at BCH were prospectively identified using a validated electronic algorithm designed to identify children with severe obesity (10), and families were approached in the clinic for recruitment in the weight management and/or primary care clinic under the Genetics of Early Childhood Obesity (GECO) study (n = 225, NCT01998750). In this cohort, 48.4% (n=109) children were of self-reported African American and/or Latina ancestry. The same algorithm was applied to the electronic data warehouse at CHOP to identify children with severe obesity retrospectively. After filtering biologically implausible data points, and removing individuals with BMI z-score < −5, or obesity duration less than 6 months, or lack of data in early childhood, 1,648 individuals with severe early onset obesity remained in the study. This was followed by analysis of longitudinal BMI trajectory by Super Imposition by Translation and Rotation (SITAR, R version 3.1.3) to generate the size, tempo, and velocity estimates for each individual subject (n=1,120) of selfreported African American ancestry (11). Children who had the highest velocity of the BMI trajectory and a DNA sample in the institutional biobank (Center for Applied Genomics) (n=157) were prioritized for genetic sequencing. At CUMC, children were recruited from the 110 families attending the Families Improving Health Together (FIT) Program, a New York state supported clinical and research program designed for children with early onset obesity (n=47).

Phenotype data for longitudinal BMI, blood pressure and laboratory studies were extracted from the electronic health records (EHRs). Class of obesity was assignedp according to the recent classification of the CDC 2000 BMI growth charts, with Class 2 as 120-140% of BMI_95ct_ and Class 3 > 140% of BMI_95pct_(9). No additional clinical studies were performed for the study.

### Molecular genetic analyses

DNA was extracted from the peripheral blood lymphocytes by standard methods or by saliva collection with DNA Genotek® kits as per manufacturer’s instructions. The coding exon (332 amino acids) and adjacent intronic regions of *MC4R* were amplified by polymerase chain reaction (PCR) using previously reported primer pairs(12). The PCR products were purified and directly sequenced using the Big Dye Sequencing kit (Applied Biosystems, Foster City, CA, USA) on an ABI 3100 automated DNA sequencer (Applied Biosystems, Foster City, CA, USA). The chromatograms from the sequencing studies were analyzed using Chromaseq software (v 1.7). A subset of the samples was sequenced by whole exome sequencing. Library preparation was performed using SureSelect XT Human All Exon V5 kit (Agilent Technologies), and sequencing was performed on a HiSeq platform (Illumina, Inc) as paired-end 2 × 125-bp runs with minimum 20X coverage. The reads were mapped to the human genome assembly (hg19; UCSC browser) using Burrows-Wheeler Alignment (BWA, 0.6.2, http://bio-bwa.sourceforge.net/). Optical and PCR duplicates were marked and removed with Picard. Local realignment of reads in the indel sites and quality recalibration were performed with the Genome Analysis Tool Kit (version 2.1, http://www.broadinstitute.org/gatk/). SNPs and small INDELs were called with GATK UnifiedGenotyper. Initial variants were filtered following GATK best practice by excluding those variants that had: 1) SNPs “QD < 2.0”, “MQ < 40.0”, “FS > 60.0”, “MQRankSum < −12.5”, “ReadPosRankSum < −8.0“; 2) Indels “QD < 2.0”, “ReadPosRankSum < −20.0”, “FS > 200.0”. ANNOVAR (version 2-1-2016, http://www.openbioinformatics.org/annovar/) was used to annotate the variants. Variants were additionally filtered to include nonsynonymous, splice site and indel variants with an allele frequency (AF) < 0.001 in Exome Aggregation Consortium (12). The variants were confirmed by Sanger sequencing. Testing was also performed in parents of 4 children with the *MC4R* variant where DNA samples were available.

### Functional studies

Selected *MC4R* variants were functionally characterized *in vitro* using cell-based assays to determine 1) the cAMP levels upon stimulation with α-MSH as a measure of MC4R signaling activity, and 2) the MC4R surface expression. Both assays were performed in HEK293 cells transiently transfected with either wild-type or mutant *MC4R*. HEK293 cells were maintained in DMEM (Cat # 11995065; ThermoFisher Scientific) supplemented with 10 % Fetal Bovine Serum (Cat # 10082147; ThermoFisher Scientific). Transfections were performed using Lipofectamine 2000 (Cat # 11668019; ThermoFisher Scientific) and Opti-MEM I (Cat # 31985062; ThermoFisher Scientific) according to manufacturer’s instructions. Wild-type (WT) *MC4R* cDNA construct (N-terminal FLAG tag, fused to EGFP at the C-terminus) was a gift from Dr. Christian Vaisse. Mutations were introduced via site-directed mutagenesis using the QuickChange Lightning Site-Directed Mutagenesis Kit (Agilent, #210518) according to the manufacturer’s recommendations. Experiments were carried out each time in duplicate and were repeated three times.

#### 1) cAMP ELISA

Cells were seeded in 24 well plates (200,000 cells in 500 μl medium per well) 20 hours prior to transfection with 0.5 μg DNA in 25 μl Opti MEM and 3 μl Lipofectamine 2000 in 25 μl Opti-MEM I per well. 24 hours after transfection the medium was aspirated, cells were rinsed once in Krebs-Ringer bicarbonate buffer containing glucose (KRBG; Cat # K4002-10X1l; Sigma-Aldrich). α-MSH (Cat # M4135; Sigma-Aldrich), de-solved in KRBG, was added at various concentrations (0, 0.001, 0.01, 0.1, 1, 10, 100 μM). Cells were incubated for 5 min at 37°C. For sample collection, the medium was aspirated, cells were rinsed once with DPBS and 200 μl 0.1 M HCl was added, followed by an incubation at RT for 10 min. Then cells were scraped off with a P1000 pipette and transferred to an Eppendorf tube and placed on ice, followed by vortexing and spinning for 15 min at 4 degrees at high speed. The supernatant was transferred into one precooled Eppendorf tubes (200 μl) and stored at −80 degrees. The ELISA was performed with the cAMP complete ELISA kit (Cat # ADI-900-163; Enzo Life Sciences) according to the manufacturer’s guidelines and 100 μl of each sample was used per reaction. For normalization, protein concentration was determined using Pierce™ BCA Protein Assay Kit (Cat # 23225; ThermoFisher Scientific).

#### 2) MC4R surface expression via surface biotinylation

Cells were seeded in 150 mm tissue culture dishes (Cat # 08-772-24; ThermoFisher Scientific) at a concentration of 16.9 × 10^6^ cells in 42 ml per dish. After 20 hours, transfection was performed with 42 μg DNA in 2111 μl Opti-MEM I and 253 μl Lipofectamine 2000 in2111 μl Opti-MEM I per dish (1 dish per experimental condition). Cells were incubated for 24 hours at 37 deg. MC4R cell surface localization was performed using the Pierce Cell Surface Protein Isolation Kit (Cat # 89881; ThermoFisher Scientific) according to the manufacturer’s instructions. Briefly, cells were rinsed once in KRBG, followed by surface biotinylation by a water-soluble and membrane-impermeable sulfo-NHS-biotin reagent and isolation of biotinylated proteins. Total proteins from each extract were loaded on a 4-12% gradient Bis-Tris gel (Cat # NP0335BOX; ThermoFisher Scientific) and transferred onto nitrocellulose membrane using iBlot 2 Dry Blotting System (ThermoFisher Scientific). Membrane was blocked for 1 hour at room temperature with 5% non-fat dry milk in TBS with 0.1% Tween-20 (Cat # 1706531; Bio-Rad) and then incubated overnight at 4°C with primary antibody against eGFP Tag (F56-6A1.2.3) from ThermoFisher Scientific (Cat # MA1-952; 1:1,000) to assess the expression of MC4R in the biotinylated cell surface fraction, washed three times using TBS with 0.1% Tween-20 and incubated with secondary antibody anti-mouse HRP (1:10,000; Cat # 7076S; Cell Signaling) for 1 hour at room temperature. Specific bands were then detected by ECL analysis using SuperSigna West Pico PLUS Chemiluminescent Substrate (Cat # 34577; ThermoFisher Scientific). Antibody against ADAM17 (Cat # ab2051; Abcam; 1:1,000), using secondary antibody anti-rabbit HRP (1:10,000; Cat # 7074S; Cell Signaling), was used as the loading control for the surface fraction.

## Results

This study included a total of 312 unrelated subjects (BCH = 109, CHOP = 157, CUMC = 46). The demographic distribution of the subjects is provided in Table 1. Briefly, 86% of the children were of self-reported African American and 20% of Latina ancestry, with 50% being female. All participating children had severe early onset obesity (see Methods).

**Table 1.** Characteristics of the study population

|  | **BCH** | **CUMC** | **CHOP** |
| --- | --- | --- | --- |
| **Subjects** | 109 | 46 | 157 |
| **Female, n (%)** | 58 (53.2) | 25 (54.3) | 74 (47) |
| **Race/Ethnicity** |  |  |  |
| Black | 56 | 5 | 157 |
| Hispanic | 67 | 42 | - |
| Black/Hispanic | 14 | 1 | NA |
| **Age at initial visit, median years (range)** | 2.25 (0.3-4.8) | 0.1 (0 – 2.1) | 0.1 (0-1.1) |
| **Age at most recent contact in years, median (range)** | 11.07 (8.3-13.1) | 9 (6.2-12.2) | 7.7 (4.8-11.4) |
| **Duration of follow-up in years, median (range)** | 5.5 (3.7-10) | 4.5 (2.3-10.5) | 6.7 (4.3-10.1) |
| **Number of measurements, median (IQR)** | 32 (35.3) | 15 (7) | 9 (7) |
| **BMI within individual over all measurements % of BMI<sub>95pct</sub>, median (IQR)** | 140 (124-155) | 135 (124-145) | 152 (135-173) |
| **Maximal BMI within individual over all measurements % of BMI<sub>95pct</sub>, median (IQR)** | 158 (146-138) | 153 (130-165) | 152 (135-173) |
| **Median Height z-score within individual over all measurements, median (IQR)** | 1.3 (0.5-1.6) | 1.3 (0.6-1.5) | 1.2 (0.5-1.8) |

BCH = Boston Children's Hospital; CUMC= Columbia University Medical Center; CHOP = Children's Hospital of Philadelphia;
%BMI<sub>95pct</sub> = percentage of 95<sup>th</sup> percentile of BMI from CDC 2000 growth charts.

Of the 312 children who underwent sequencing of *MC4R*, eight individuals were heterozygous for rare *MC4R* nonsynonymous protein coding variants with minor allele frequency (MAF) < 1 %. The variants identified were R7S, F202L (n=2), M215I, G252D, V253I, I269N, and F284I, three of which have not been previously described in the literature (M215I, G252D, F284I). A further fifteen children harbored common (MAF>1%) synonymous or nonsynonymous coding variants (V103I, I198=, Q156=, I251L).

The BMI trajectories of the children carrying these variants did not differ from the rest of the cohort (Figure 1 a), and there was no increase in the height SD when compared with those of comparable obesity without the mutation, unlike prior report (13) (Figure 1 b). The clinical features of the subjects are summarized in Table 2. There was no history of consanguinity in any of the families (Fig. 1, c-h). In the 4 families where parental DNA was available, autosomal dominant inheritance was present in 3 (Fig. 1c, d and g), while 1 child had a *de novo* mutation (Fig.1 e).

**Figure 1.**
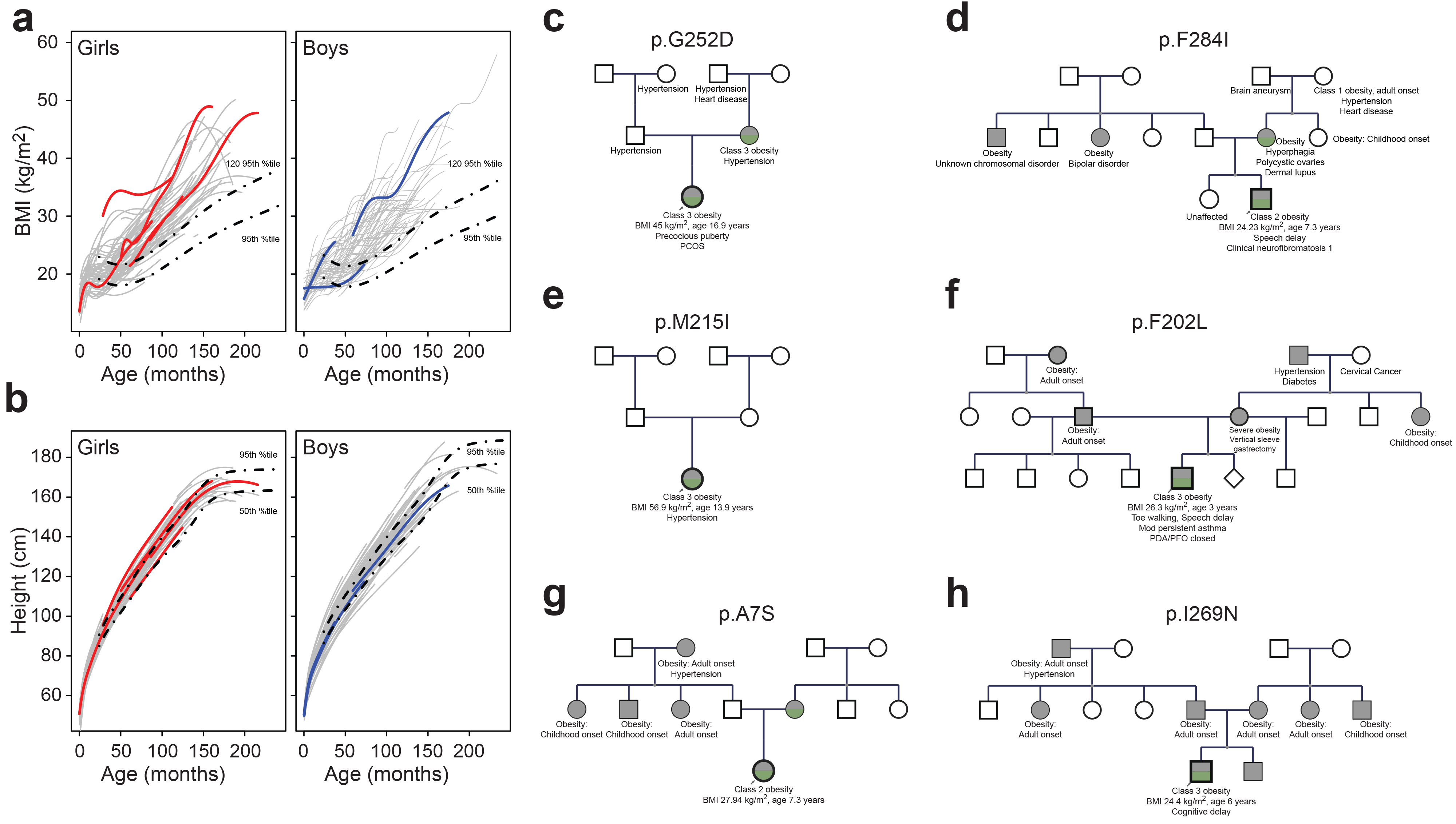
a: BMI trajectories fitted with SITAR model of the children enrolled in the study. The colored lines represent children with mutations in MC4R and grey lines are those without. The dashed lines represent the BMI curves based on CDC 2000 growth charts. b: Height trajectories fitted with SITAR model of the children enrolled in the study. The colored lines represent children with mutation in MC4R and grey lines are those without. The dashed lines represent the height curves based on CDC 2000 growth charts. c-h: Pedigrees of 6 subjects with mutations in MC4R where family data was available. Grey filled figures are individuals with obesity. Green color represents documented mutation in MC4R gene.

**Table 2.** Phenotype and variant details of individuals with *MC4R* mutations

| Phenotype details |  |  |  |  |  |  |  |  |
| --- | --- | --- | --- | --- | --- | --- | --- | --- |
| Amino acid change | p. G252S | p. F284I | p.R7S | p.M215I | p. V253I | p. F202L | p. I269N | p.F202L |
| c.DNA change | c.755 G>A | c.850 T>A | c.19C>T | c.645G>A | c.757 G>A | c.606 C>A | c.806 T>A | c.606 C>A |
| Center | BCH |  |  |  | CHOP |  | CUMC |  |
| Race/Ethnicity | AA | Latino | AA | Latino | AA | AA | Latino | Latino |
| Gender | F | M | F | F | F | F | M | M |
| Birth weight (kg) |  |  |  |  |  |  | 3.93 | 3.375 |
| Obesity Class | 3 | 2 | 2 | 3 | 2 | 3 | 3 | 3 |
| First available age (months) | 48 | 24 | 28 | 37 | 36 | 81 | 0 | 0 |
| First available BMI (%BMI<sub>95pct</sub>) | 153.1 | 119.5 | 119.7 | 133.3 | 165 | 124 | 90 | 50 |
| Last available age (months) | 203 | 87.7 | 87.5 | 160 | 99 | 123 | 80 | 33.5 |
| Last available Height (cm) | 169.5 | 125.2 | 133.5 | 167.3 | 154 | 145.5 | 119 | 92.7 |
| Last available Height (%tile) | 84.6 | 60.65 | 95.7 | 86.8 | >99 | 80 | 56 | 40 |
| Last availableBMI (%BMI<sub>95pct</sub>) | 152.3 | 126.6 | 140.3 | 200.6 | 132 | 174 | 130 | 148 |
| Last available BMI (kg/m2) | 45.01 | 24.23 | 27.94 | 56.9 | 34.87 | 33.4 | 22.3 | 26.3 |
| BP |  |  |  |  |  |  |  |  |
| Age (months) | 215 | 99 | 87 | 160 | 98 | 123 | 71 | 33.5 |
| Height (cm) | 169 | 130 | 134 | 167.3 | 152 | 145.5 | 116 | 92.7 |
| SBP (%tile) | 130 (94) | 99 (48) | 98 (40) | 127 (92) | 104(53) | 129 (99) | 97 (59) | 95 (72) |
| DBP (%tile) | 67 (50) | 50 (20) | 72 (86) | 65 (44) | 56(24) | 60(45) | 56 (65) | 49 (67) |
| Variant Details |  |  |  |  |  |  |  |  |
| gnomAD Total Allele Frequency | 4.95E-05 | Absent | 4E-06 | 8E-06 | 6E-05 | 0.0009 | 0.00092 | 0.000898 |
| gnomAD Allele Frequency Africans | 0 | 0 | 0 | 0 | 0 | 0.00942 | 0 | 0.009415 |
| gnomAD Allele Frequency Latino | 0.00011 | 0 | 0 | 0 | 0.00011 | 0.00045 | 0.00731 | 0.000452 |
| Polyphen | Probably damaging | Probably damaging | Benign | Probably damaging | Probably damaging | Benign | Probably damaging | Benign |
| SIFT | deleterious | deleterious | tolerated | deleterious | deleterious | tolerated | deleterious | tolerated |
| CADD score | 28.6 | 26 | 12.25 | 28.7 | 25 | 12.17 | 26.8 | 12.17 |
| Functional | known | Novel | Novel | known | known | known | known | known |
| References | Hinney A, 1999, 2003, 2006; Lubrano-Berthelie C, 2003, 2006; Xiang Z, 2006; Stutzmann F, 2008 |  |  | unpublished | Yeo GS, 2003; Farooqi S, 2003; Nijenhuis WA, 2003; Lubrano-Berthelie C, 2003 | Tao YX, 2005; Stutzmann F, 2008; Hughes DA, 2009; Calton MA, 2009; Xiang Z, 2010; Hohenadel MG, 2014 | Tan K, 2009; Calton MA, 2009; Thearle MS, 2012; Hohenadel MG, 2014; | Tao YX, 2005; Stutzmann F, 2008; Hughes DA, 2009; Calton MA, 2009; Xiang Z, 2010; Hohenadel MG, 2014 |

AA = African American; F = female; M = male; NA = not available; TG= Triglycerides
Class 1 obesity = BMI 95-120% of BMI<sub>95pct</sub>; Class 2 obesity= BMI 120-140% of BMI<sub>95pct</sub>; Class 3 obesity = BMI > 140% of BMI<sub>95pct</sub>

A subset of the study subjects had data for cardiometabolic risk factors: blood pressure (n = 140), hemoglobin A1c (n = 121), lipid panel (n = 111), and liver enzymes (n = 122). The proportion of individuals with diabetes was higher in the subjects with *MC4R* mutation (12.5% vs 0.8%, p = 0.01), and prediabetes was not different (25% vs 15%, p = 0.45); while the abnormalities in lipid panel and liver enzymes were not different. Blood pressure in the *MC4R* positive subjects was not lower as previously reported(14); on the contrary, the systolic BP > 95^th^ percentile for age, gender and height was noted in 2 individuals (Table 2). The proportion of children with age of onset of severe obesity < 2 years was higher in those with *MC4R* mutation (75% vs 43%, p= 0.07).

### Clinical Features of Subjects

#### p.G252S, 18:58038828 C>T, c.755 G> A

This 17-year old female African-American girl with history of severe obesity since 9 months of age was first seen in the medical system to rule out precocious puberty at 4-4/12 years of age with Class 3 obesity (BMI z= 3.45, 153% of BMI_95pct_) that continued at 17 years of age. Her linear growth tracked along the 90^th^ percentile till 12 years of age at which time it reached a plateau with the onset of puberty with adult height in the 80^th^ percentile. Her mother has a history of obesity, carries the same *MC4R* variant, and the maternal family has history of cardiometabolic diseases. The patient is currently being treated for polycystic ovary syndrome, severe obesity and prediabetes. The variant, G252S, is absent in ClinVar and reported primarily in Latino (AF_Lat_ 0.0001) individuals in gnomAD (15).

#### p.F284I, 18:58038733 A> T, c.850 T>A

This 8-year-old boy of Puerto Rican descent had a birth weight of 4 kg. He was first seen at the hospital at 2 years of age with BMI of 22.2 kg /m2 (BMI z= 3.37, 120% of BMI_95pct_). He was diagnosed with speech delay that resolved with therapy and cafe-au-lait spots that have grown with age. He does not have any other signs of neurofibromatosis-1, and negative for mutation in *NF-1* gene. His BMI continues to be in Class 2 obesity category with nutritional/psychological intervention, the most recent BMI being 26.60 kg/m2 (BMI z = 2.39, 127% of BMI_95pct_). His height is currently within the 50-60^th^ percentile. Both parents are obese, with high cardiometabolic morbidity in the maternal family. The variant is absent in ClinVar, gnomAD and HGMD.

#### p.R7S, 18:58039564 G>T, c.19C>T

This is a 7-year old African American girl, who started gaining weight at 22 months of life. Her BMI was 22.50 kg/m2 (BMI z= 3.27, 120% of BMI_95pct_) at 2 years of age that has consistently remained in Class 2-3 obesity with the current BMI at 27.90 kg/m2 at 7.25 years of age (BMI z = 2.66, 140% of BMI_95pct_). Her height was in 72th percentile in early childhood and at 95^th^ percentile at 7 years. There is extensive family history of Type 2 diabetes, but the child does not have any cardiometabolic abnormalities. The *MC4R* variant is inherited from her mother. The variant is absent in ClinVar and reported in 1/248964 alleles in gnomAD in an individual of South Asian inheritance (AF_SEA_ 0.00003).

#### p.M215I. 18:58038938 C>T, c.645G>A

This 14-year-old girl of Latino ancestry had a birth weight of 3.5 kg. The family reports history of obesity since 8 months of age, and the first recorded BMI at 3 years is 24.30 kg/m2 (BMI z = 3.60, 132% of BMI_95pct_). Her height has consistently been in the 97^th^ percentile, and BMI at 14 years is 57.50 kg/m2 (BMI z= 2.91, > 200% of BMI_95pct_). At 14 years of age, she has hypertension treated with Lisinopril, dyslipidemia, prediabetes with an elevated HbA1c and impaired fasting glucose, hyperinsulinemia and extensive acanthosis. The variant is absent in ClinVar, and reported in 2/251254 heterozygous alleles in individuals of Non-Finnish European descent in gnomAD (AF_NFE_ 0.000008) only.

#### p.V253I, 18: 58038826 C>T, c.757 G>A

This is an 8.5 years old African-American girl with severe obesity since 2.3 years (BMI z = 4.7, 174% of BMI_95pct_). The available growth data demonstrate tall stature (height Z-score 1.80 – 2.37 between ages 2.3 – 3.1 y). At study intake, the only noted medical problem was obesity, and at age 7, she was being treated for type 2 diabetes mellitus requiring insulin. This variant has been reported as pathogenic in heterozygous state in ClinVar and 17/282,740 alleles in gnomAD, primarily in Latino (AF_Lat_ 0.0001).

#### p.F202L, 18: 58038977 G>T, c.606 C>A

The subject is a 10-year old African-American female had an increase in BMI after 2 years of age to a maximal observed at 7 years (BMI z =2.63, 132% of BMI_95pct_). She is noted to be shorter than typical at younger ages (height z = −1.77 at age 3.2 years) that changed to average (height z = −0.41) at age 7 years. Aside from obesity, the only documented co-morbidity was asthma. This variant has been reported with conflicting interpretations of pathogenicity and seems to be limited to individuals of African American and Latino ancestry in gnomAD Allele Count (AC) 254/282,718 alleles of which 235 were reported in African (AF_Afr_ 0.0094, 1 homozygote), and in Latino individuals (AF_Lat_ 0.00045), and none in Europeans.

#### p.I269N, 18:58038777 A> T, c.806 T> A

This child is a 6-year-old boy of Latino ancestry, with BMI of 18.1 kg/m2 at 16 months of age (Class 1 obesity). At 5 years of age, his BMI is 24.4 kg/m2 (BMI z = 3.4, 135% of BMI_95pct_). His height has been between 34–60^th^ percentile, and aside from acanthosis, there are no other cardiometabolic risks secondary to obesity. He was diagnosed with speech delay that has resolved, but continues to receive therapy for cognitive delay and treatment for management of obesity. This variant has been reported with conflicting evidence of pathogenicity in ClinVar and 260/282764 alleles (AF_total_ 0.0009195) in gnomAD with 259 alleles in individuals of Latino descent (AF_Lat_ 0.0073, 5 homozygotes), and none in Europeans or African.

#### p.F202L, 18: 58038977 G>T, c.606 C>A

This 3-year old Latino boy was born to a mother with severe obesity (reported BMI 40 kg/m^2^). Pregnancy was complicated by maternal pre-eclampsia, polyhydramnios and gestational diabetes and the baby was delivered at 34 weeks by cesarean section with a birth weight of 3.4 kg and macrosomia. The child was diagnosed with gastrointestinal reflux, moderate persistent asthma and toe walking. Weight gain was noted at 12 months with BMI of 19.45 kg/m2 and has continued at 3 years of age with BMI 26.3 kg/m2 (BMI z= 4.69, 148% of BMI_95pct_). Height has been along the 50^th^ %tile for age. Karyotype and chromosomal microarray are normal with normal DNA methylation studies for Prader Willi and Angelman syndrome. Both parents are obese and mother has undergone vertical sleeve gastrectomy after the birth of the child.

### Functional follow-up

To understand the clinical relevance of the identified rare variants, we performed functional assays on four variants, one novel (F284I) one previously characterized but as yet unpublished (M215I) (www.mc4r.uk.org), and one previously characterized and two published V253I and G252S (16). As the primary signaling pathway for MC4R involves the activation of Gα_s_ subunit and, consequently, adenylyl cyclase (17–19), we measured the half-maximal effective concentration (EC_50_) of ligand-induced cyclic AMP (cAMP) in transiently transfected HEK293 cells expressing wild-type or mutant *MC4R* after exposure to a range of concentrations of endogenous ligand α-MSH (Figure 2 a). In this experiment, the initiation in activating Ga_s_ signaling required 10-fold higher α-MSH concentration in 2 variants -- M215I (EC_50_ ratio 3.9) and G252S (EC_50_ ratio 5.9) -- and 100-fold higher in the F284I variant (EC_50_ ratio 20.9) compared to the wild-type (WT) MC4R. Of note, none of the tested variants produced a peak cAMP response as high as that achieved with WT MC4R protein, suggesting that even the variants with normal EC_50_ ratios have a functional defect. Cell surface localization of the receptor in MC4R-expressing HEK293 cells was assessed by biotinylation of MC4R (Figure 2 b, See Methods). Western blot analysis with antibodies against EGFP to detect the MC4R-EGFP fusion protein showed diminished levels for the mutants as compared to WT, suggesting that decreased expression or protein half-life could contribute to the observed *in vitro* functional defects for all of these variants. For the variant V253I, while the dose response curve was close to normal, the surface expression was nearly absent, emphasizing the defect in surface trafficking.

**Figure 2.**
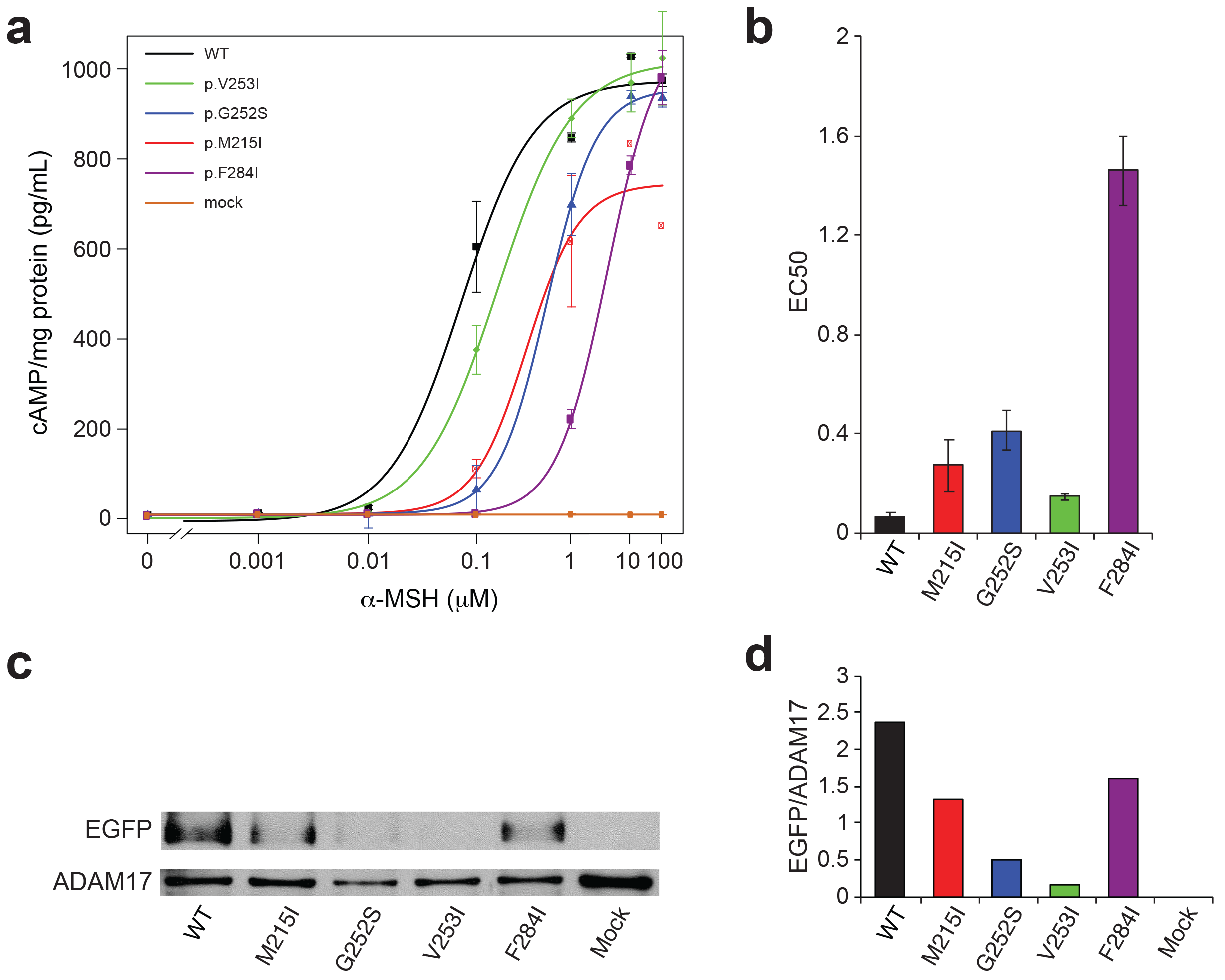
a: Dose response curves (DRC) for the tested MC4R mutants along with WT MC4R as the positive control and mock as the negative control in HEK-293 cellular overexpression system. The DRCs are shifted to the right for all the mutants, and statistically different in all tested variants except V253I. b: EC50 for optimal cAMP response measured by ELISA. All variants except V253I have p-value < 0.05 when compared to WT. c: Western blot gel of the surface expression of MC4R and ADAM17 protein in HEK-293 cells measured by surface biotinylation. All mutants have lower expression of the protein compared to WT. d: Quantitative measurement of the ratio of EGFP/ADAM17 protein expression using ImageJ software for surface expression of the protein. All mutants were significantly lower than WT.

## Discussion

Mutations in the leptin-melanocortin pathway are the best-defined causes of monogenic obesity, of which autosomal dominant mutations in *MC4R* are the most common. While the prevalence of severe obesity is highest in children from non-European backgrounds, there is little information on the monogenic causes in these populations.

To our knowledge, this cohort of 312 children with severe early onset obesity from selfreported African American and/or Latina ancestry is the largest reported cohort of severely obese children from these populations tested for *MC4R* variants. Rare pathogenic *MC4R* variants, shown either by functional testing or previous literature reports, were found in eight (2.6%) children. This prevalence is comparable to that reported in recent studies in children of European ancestry (4,20), but lower than that reported by Farooqi *et al*(5). There are two prior reports of rare variants in *MC4R* in individuals of African American descent: in a cohort of 25 children from the Guadeloupe Island in the Caribbean, one 11.8 years old boy with severe obesity (BMI 30.5 kg/m2, BMI z= 2.39), insulin resistance and blood lipid abnormalities was found to have a variant in *MC4R*, I301T that is absent in gnomAD and ClinVar (7); in a report of 167 adults of African descent in South Africa, Logan *et al* identified five rare variants (R7H, R165Q, I170V, I198T, F202L, I251V). In gnomAD, low frequency variants in *MC4R* like I251L and V103I are equally distributed across populations of different ancestries, while some rare variants are more frequently seen in individuals of African descent such as N240S (AF_Afr_ 0.002), F202L (AF_Afr_ 0.009), R7H (AF_Afr_ 0.0004); or Latino L304F (AF_Lat_ 0.0003), T276C (AF_Lat_ 0.0004), Ile269Asn (AF_Lat_ 0.0073), A259V (AF_Lat_ 0.0002), T150I (AF_Lat_ 0.0005) or East Asian such as M218T (AF_EastAs_ 0.0005), Y35C (AF_EastAs_ 0.001). These data and our results suggest that the repertoire and distribution of rare functional variants in *MC4R* in obese individuals of African American and/or Latina ancesty differ from those seen in the studies of individuals of European ancestry.

Our sample is enriched for severe early onset obesity (BMI > 120% of _BMI95pct_ with an onset of obesity prior to 6 years of age), and the onset of obesity in most children with the *MC4R* mutations was prior to 2 years of age, similar to other recent reports (21,22). We did not observe any differences in the BMI trajectories of the children with the mutation in our cohort. This observation is in keeping with the finding by Kohldorf *et al*, where they observed a much higher BMI at an early age in children with *LEP* and *LEPR* mutations, as compared to those with *MC4R* (21). Additionally, we did not observe differences in stature of the individuals with or without the mutations. It is possible that this phenotype is more likely to be observed later in life, or is different across ancestries, or that we are insufficiently powered to observe small differences in height.

Overall, *the MC4R* gene has been extensively studied, and the recently curated atlas of human *MC4R* variants (www.mc4r.uk.org) has provided a valuable resource to review the functional relevance of rare variants. In this study, three out of eight (38%) of the variants that we observed (M215I, G252D, F284I) had not been previously described in the literature or in publicly available clinical resources for genetic variants (www.clinvar.org), likely reflecting both increased genetic diversity in children across different ancestry/ethnicity as well as substantial allelic heterogeneity for pathogenic variants in this gene.

In this cohort of severe early onset obesity, none of the enrolled patients had undergone clinical genetic testing for *MC4R* despite their extreme phenotype seen at an early age. With the advent of safer melanocortin agonist therapies that have been used in patients with *POMC*, *LEPR* and *MC4R* mutations (23,24), and the increasing focus on personalized medicine, clinical genetic testing of children with severe early onset obesity is becoming increasingly relevant. Studies on the relationship between genetic and phenotypic variation have been historically carried out on people of European ancestry, with much lower representation of other ancestries/ethnicities. Appropriately, attention is increasingly being paid to the valuable insights that can be gleaned from studies in genetically diverse populations. The clinical role for genetic testing in severe, early onset obesity has been discussed in the guidelines for the management of obesity in children by the Endocrine Society (25), and we agree that, where indicated, opportunities for testing be made available across the wide range of affected children. Specifically, in order to interpret genetic testing in the populations most affected, it will be critical to have allele frequency and functional data on variants from children of diverse backgrounds.

## Acknowledgements

We gratefully acknowledge the support and critical review of the manuscript by Drs. Rudolph Leibel and Wendy Chung from Columbia University Medical Center. This work is supported in part by the Yale and Baylor-Hopkins Center for Mendelian Genomics. Clinical Translational Study Unit at Boston Children’s Hospital and Nutrition and Obesity Research Center at Harvard. The Yale Center for Mendelian Genomics (NIH M#UM1HG006504-05) is funded by the National Human Genome Research Institute and the National Heart, Lung, and Blood Institute. The GSP Coordinating Center (U24 HG008956) contributed to cross-program scientific initiatives and provided logistical and general study coordination. The Baylor-Hopkins Center for Mendelian Genomics if funded through National Human Genome Research Institute grant 5U54HG006542. (BCH CTSA grant) and Nutrition and Obesity Research Center at Harvard if funded by NIH-P30 DK040561.

